# Multiplex Genome Editing Overcomes Photoperiod Sensitivity in Tropical Maize

**DOI:** 10.64898/2026.08.05.742845

**Authors:** Keunsub Lee, Ella Hampson, Rina Carrillo, Minjeong Kang, Fernanda Ghenov, Lauren Higa, Zhi-Yan Du, Gregory R. Schoenbaum, Jianming Yu, Kan Wang, Michael G. Muszynski

**Affiliations:** Department of Agronomy, Iowa State University, Ames, Iowa, USA; Crop Bioengineering Center, Iowa State University, Ames, Iowa, USA; Department of Tropical Plant and Soil Sciences, University of Hawaii at Manoa, Honolulu, Hawaii, USA; Interdepartmental Plant Biology, Iowa State University, Ames, Iowa, USA; Department of Molecular Biosciences & Bioengineering, University of Hawaii at Manoa, Honolulu, Hawaii, USA

## Abstract

Tropical maize is a rich source of genetic diversity that could enhance temperate maize breeding programs, but its sensitivity to long-day photoperiods, resulting in delayed flowering, limits its widespread use. To overcome this barrier, the Genome Engineering to Sustain Crop Improvement (GETSCI) project used CRISPR/Cas9 to mutate three flowering repressor genes, *ZmCCT9, ZmCCT10*, and *ZmRAP2*.7, in the tropical inbred Tzi8. A single sgRNA targeting the first exon of each target gene was combined with an excision cassette carrying the morphogenic genes *Babyboom* (*Bbm*) and *Wuschel2* (*Wus2*) to enable efficient transgenic plant regeneration. Transgenic plants carrying frameshift edits in each target gene were recovered, and subsequent crosses produced two non-transgenic genotypes: a double-edited *zmcct10, zmrap2*.*7* line and a triple-edited *zmcct9, zmcct10, zmrap2*.*7* line. Multiple flowering traits were measured for the edited genotypes and unedited Tzi8 inbred in short-day (Hawai‘i) and long-day (Iowa) field conditions. Both edited genotypes flowered significantly earlier than Tzi8 in both environments. Notably, under long-day conditions, flowering of the two edited lines overlapped with that of the temperate inbred B73, whereas Tzi8 did not. Together, these results demonstrate that targeted, multiplex gene editing can reduce photoperiod sensitivity in a tropical inbred, expanding access to previously untapped genetic diversity for temperate maize improvement.

---

To Editor,

Maize (*Zea mays* L.) improvement depends on access to genetic diversity, but its use is limited by barriers to hybridizing diverse germplasm. During maize expansion from tropical to temperate regions, selection for earlier flowering and reduced photoperiod sensitivity enabled adaptation to long-day environments. Most tropical germplasm, however, remains photoperiod sensitive and flowers much later—or fails to flower—under long days, creating asynchronous flowering that restricts the incorporation of valuable tropical diversity into temperate breeding programs (Choquette et al. 2023).

Flowering time is regulated by interacting aging and photoperiod pathways that converge on the floral activator *ZEA CENTRORADIALIS8 (ZCN8)* (Fig. 1a; Stephenson et al. 2019). In the aging pathway, the *APETALA2-like* transcription factor *ZmRap2*.*7* represses *ZCN8* expression (Liang et al. 2019), while in the photoperiod pathway the duplicated genes *ZmCCT9* and *ZmCCT10* suppress *ZCN8* under long-day conditions (Hung et al. 2012; Yang et al. 2013; Huang et al. 2018). Evolutionary changes affecting these pathways facilitated maize adaptation from tropical to temperate environments.

**Figure 1.**
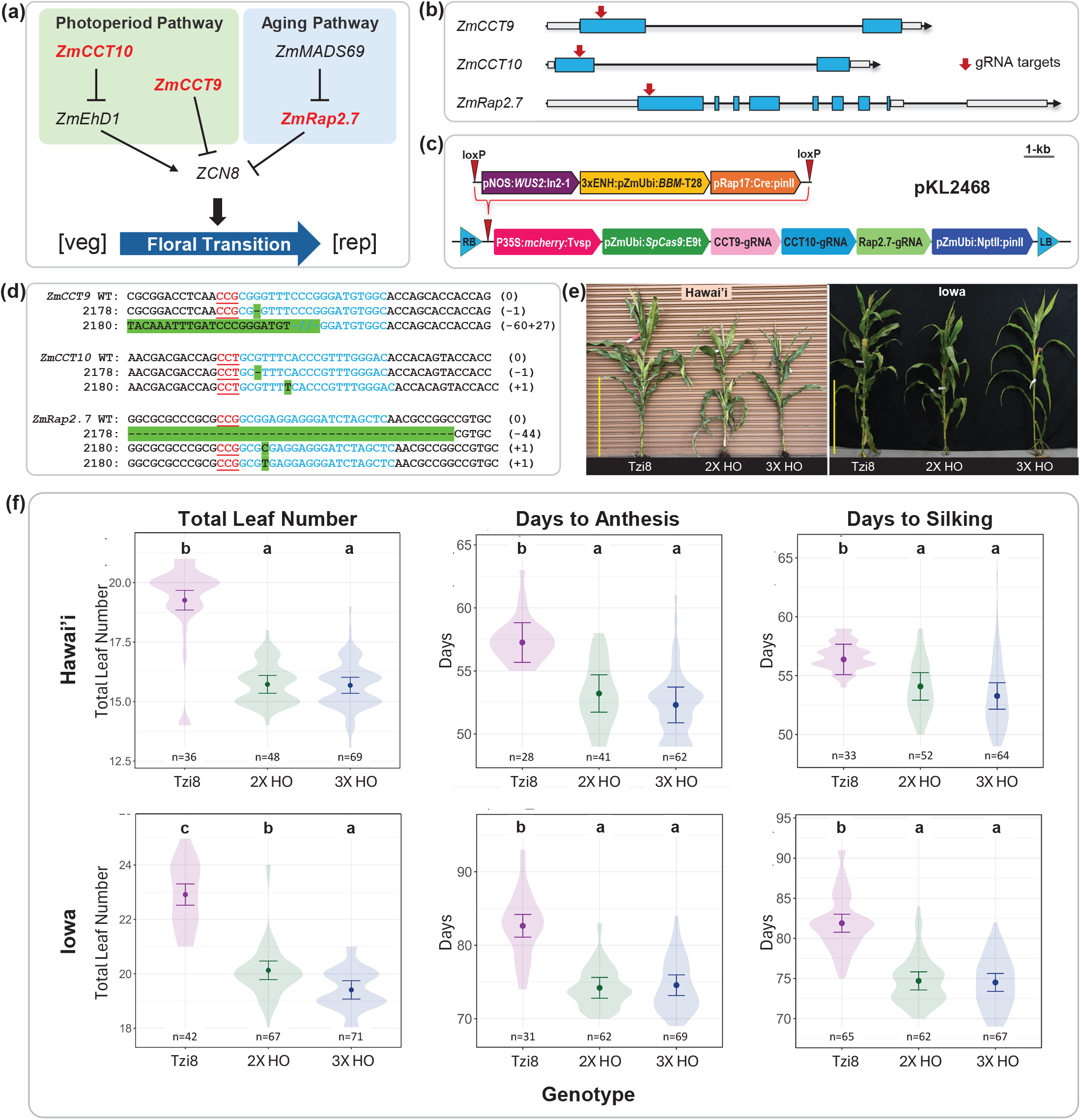
Engineering photoperiod insensitive tropical maize. (a) Simplified flowering-time network showing edited repressor genes (red) that suppress ZCN8, a promoter of vegetative [veg]-to-reproductive [rep] transition. (b) Gene models of ZmCCT9, ZmCCT10, and ZmRap2.7 with gRNA target sites (red arrows). (c) CRISPR/Cas9 construct used for Tzi8 transformation. (d) Mutations in T1 plants #2178 and #2180 that generated fixed edited lines; sgRNA targets are blue, PAMs red-underlined, and edits green. (e) Flowering phenotypes of Tzi8, 2X HO, and 3X HO lines in Hawai’i (short days) and Iowa (long days). Yellow bar = 1 m. (f) Flowering traits under both photoperiods; violin plots show distributions, estimated marginal means ±95% CI, and significant genotype differences (adjusted P < 0.05). *n*, number of plants.

Building on this knowledge, we used multiplex CRISPR/Cas9 genome editing to simultaneously disrupt *ZmRap2*.*7, ZmCCT9*, and *ZmCCT10*. The tropical maize line Tzi8 was used for *Agrobacterium*-mediated immature embryo transformation (Masters et al. 2020; Supporting Information). To induce edits, three single-guide RNAs (sgRNAs) were designed to target the first exons of *ZmCCT9, ZmCCT10*, and *ZmRap2*.*7*, respectively (Figure 1b). To address the low regenerative capacity of Tzi8 embryonic calli, the binary vector pKL2468 included the morphogenic genes *Bbm* and *Wus2* and the abscisic acid-induced *Cre* recombinase cassette flanked by two *lox*P sites. This newly developed vector also carried a red fluorescent *mCherry* marker, the maize codon-optimized *SpCas9* cassette, the three sgRNAs, and the plant selectable marker *neomycin phosphotransferase II* gene (*NptII*) (Figure 1c; Supporting Information; Kumar et al. 2025).

From a total of 412 *Agrobacterium-*infected immature embryos, we obtained ten regenerated plants: five transgenic events with successful morphogenic gene excision, four non-excised events with developmental abnormalities, and one non-transgenic escape (Table S1). Interestingly, all nine T0 events were obtained from the same ear (Ear 2), suggesting that immature embryo quality is an important factor for successful regeneration in Tzi8. These five transgenic events were acclimatized to short-day (Hawai’i) greenhouse conditions and grown to maturity.

To verify gene editing, target gene regions were amplified by PCR (Table S2), and amplicons were analyzed using Sanger sequencing, either directly or after subcloning. Sequence trace files were analyzed using Tracking of Indels by Decomposition (TIDE) and Inference of CRISPR Edits (ICE) analyses (Supporting Information). The five T0 events each carried indel mutations at the three target genes with varying efficiencies. Of these, three events (events #1038, #1039, and #1042) produced T1 seeds after crossing with the wildtype Tzi8 inbred (Table S3).

T0 event #1042 was prioritized for further analysis, as it exhibited high knock-out scores for all three genes (Table S3). After outcrossing this event to Tzi8, its T1 progeny were genotyped to identify plants carrying the transgene and parental or new indel mutations in all three target genes (Figure 1d, Tables S4 and S5). The selected plants were either sib-pollinated or back-crossed to Tzi8 to generate transgene-free T2 progeny (Figure S1, Table S6). PCR genotyping identified several transgene-free T2 plants, of which, #1330-5 (Table S6), was identified as heterozygous edited at *ZmCCT9* and biallelic edited at *ZmCCT10* and *ZmRap2*.*7* (Figure S1). To produce fixed edited lines, T2 plant #1330-5 was self-pollinated and all its T3 progeny (∼ 190 plants) were genotyped using a combination of PCR amplification followed by Sanger sequencing; cleaved amplified polymorphic sequences (CAPS) marker analysis for the *zmcct9* and *zmrap2*.*7* deletion alleles; and PCR-amplicon cloning and clone sequencing (Methods S1; Figure S1, Table S6). Among the T3 progeny, we obtained unambiguous genotypes for 27 double-homozygous (2X HO; *zmcct10* and *zmrap2*.*7*) and 33 triple-homozygous (3X HO; *zmcct9, zmcct10*, and *zmrap2*.*7*) edited plants (Figure S1, Table S6). Self-pollinating these homozygous T3 plants created fixed double- and triple-edited T4 lines (Figure S1), which were evaluated together with the Tzi8 inbred for flowering phenotypes in the summer 2025 field under short-day (Hawai’i, max. daylength 13.5 hrs.) and long-day (Iowa, max. daylength 15.25 hrs.) conditions (Methods S1).

Both the double- and triple-edited lines flowered significantly earlier than Tzi8 under both field conditions, based on pairwise comparisons of estimated marginal means (EMMs) from the linear mixed-effects model (Methods S1, Figures 1e, f). In both Hawai’i and Iowa, the edited lines produced 2-4 fewer leaves than Tzi8 (Supporting Data Tables S7 and S8) consistent with an accelerated floral transition. Analysis of shoot apical meristem development further confirmed an earlier floral transition for the double- and triple-edited plants compared with Tzi8 in short-day conditions (Figure S2). Under long-day conditions, the triple-edited line produced significantly fewer leaves than the double-edited line (EMMs: 19.4 vs 20.1, adj. *P* < 0.001), supporting a role for *ZmCCT9* in long-day-dependent flowering repression (Table S8, Figures 1e, f). Anthesis and silking also occurred significantly earlier in both edited lines than in Tzi8 (Figure 1f), with anthesis accelerated by ∼4-5 days in Hawai’i and ∼8 days in Iowa (adj. *P* < 0.001; Tables S9 and S10). Importantly, flowering of these edited tropical lines was now synchronized with that of several temperate inbreds under long-day conditions (Figure S3), effectively circumventing a major reproductive barrier. These near-isogenic edited lines, preserving the entire tropical Tzi8 genome, provide a faster and more precise conversion than the conventional time-consuming backcrossing breeding.

Multiplex CRISPR/Cas9 editing of three flowering repressors generated transgene-free tropical maize that flowers under long-day temperate conditions while retaining its tropical genetic background. Combined with Bbm/Wus2-enabled transformation, this provides an efficient genome-engineering pipeline for a transformation-recalcitrant genotype and produces breeding germplasm with reduced photoperiod sensitivity. Beyond defining gene function, this work delivers a practical breeding technology and a framework for engineering photoperiod adaptation in maize and other crops.

## Supporting information

All Supporting Information

## Author contributions

MGM, KW, and JY conceived the project. LH, ZD, KL, MK, FG, EH, RC and GS performed the experiments and analyzed results. KL, MGM, and KW drafted the manuscript, which all authors reviewed.

## Acknowledgments

This project was partially supported by National Science Foundation (NSF) Plant Genome Research Program award IOS-1917138 to KW and KL, by the NSF Established Program to Stimulate Competitive Research’s Research Infrastructure Improvement Program award 2121410 to MGM, ZD, KW, KL, and JY, by the seed grant fund from Crop Bioengineering Center of ISU to KL, by the National Institute of Food and Agriculture of United State Department of Agriculture Hatch project #IOW05768, and by State of Iowa funds. KW’s contribution to this work is partially supported by (while serving at) the National Science Foundation.

## Data Availability Statement

The data that support the findings of this study is available in the Supporting Information of this article.

## Notes

### Competing Interest Statement

The authors have declared no competing interest.

