## Supplementary material for "Multiplex Genome Editing Overcomes Photoperiod Sensitivity in Tropical Maize": All Supporting Information

Brief Communication

**Methods**

1. CRISPR/Cas9 vector construction

   1.1 Gateway Destination Vector pKL2444 construction

   A Gateway destination vector pKL2444 (27,543 bp) was constructed based on the binary vector pKL2393 (Zobrist et al. 2023), which has the pVS1 origin of replication (ori) for *Agrobacterium*, pBR322 ori for *E. coli*, the kanamycin resistance gene for bacterial selection, *mCherry* as a visual marker, and *neomycin phosphotransferase II* (*NptII*) for plant selection. Two pairs of PCR primers (Table S2) were used to amplify the 11.3 kb fragment containing the morphogenic genes *Babyboom* (*Bbm*) and *Wuschel2* (*Wus2*), and *Cre* recombinase expression cassettes from the PHP101515 construct (Kumar et al. 2025). Amplified morphogenic gene fragments were assembled with *Pme*I digested pKL2393 DNA using the NEBuilder® HiFi DNA assembly master mix (NEB, MA, USA) according to the manufacturer's instructions. The resulting 27.5 kb destination vector pKL2444 was used for the 3-way Gateway reactions to clone *SpCas9* and three sgRNA expression cassettes.

   1.2 *ZmCCT9*, *ZmCCT10*, *ZmRap2.7* gRNA design

   We used three sgRNAs to individually target *ZmCCT9*, *ZmCCT10*, and *ZmRap2.7*, respectively. CHOPCHOP v3 (Labun et al. 2019) was used to design these sgRNAs and their editing efficiency was evaluated using the maize protoplast transient transfection method (Higa et al. 2025). We chose ZmCCT9-gRNA1, ZmCCT10-gRNA1, and ZmRap2.7-gRNA1, which demonstrated highly efficient targeted editing in Tzi8 protoplasts (Higa et al. 2025).

1.3 pKL2468 construct assembly

Selected sgRNA spacer sequences (Table S2) were individually cloned into *Esp*3I digested entry vectors as previously described (Lee et al. 2023). Three sgRNA cassettes were assembled into the Gateway entry vector pYPQ143 via Golden Gate cloning using *Bsa*I and T4 DNA ligase (Lowder et al. 2015). For final assembly, Gateway® LR clonase enzyme mix II (ThermoFisher, MA, USA) was used for a 3-way Gateway reaction that included the Destination vector pKL2444, maize codon optimized *SpCas9* entry vector pYPQ166 (Lee et al. 2019), and the pYPQ143 with three sgRNAs. The resulting 32,646 bp construct was named pKL2468.

2. *Agrobacterium*-mediated Tzi8 immature embryo transformation

2.1 *Agrobacterium* transformation

The T-DNA construct pKL2468 was introduced into a *recA*-deficient *Agrobacterium* strain LTR1A (Azanu et al. 2026), which was derived from the thymidine auxotrophic strain LBA4404Thy- (Ranch et al. 2012) and harbored a virulence gene helper plasmid pKL2299A (Aliu et al. 2024). This was because the maize *Ubiquitin* promoter was used for multiple gene cassettes and spontaneous recombination can lead to the loss of a major portion of the T-DNA region. Colonies resistant to gentamicin (50 mg/L) and kanamycin (50 mg/L) were grown in liquid culture and glycerol stocks were prepared.

2.2 Tzi8 immature embryo transformation

We used a modified QuickCorn method as previously reported (Masters et al. 2020). Briefly, Tzi8 ears, grown in the University of Hawai'i greenhouse, were harvested 12-14 days post-pollination, packed in a cooler with cold packs, and shipped, by FedEx next day, to Iowa State University. Upon arrival 1.8-2.5 mm immature embryos were isolated after surface disinfection. After 5-min infection with *Agrobacterium* cell suspension ($OD_{550}$ = 0.45 – 0.55), immature embryos were placed on cocultivation medium with scutellum side up, and co-cultured at 20˚C overnight (16-20 h) in the dark. The next day, embryos were transferred to resting medium and incubated at 28˚C in the dark for 5 days. After the resting period, growing coleoptiles were removed and the embryos were placed on a maturation medium supplemented with 50 µM abscisic acid and incubated at 28˚C in the dark for 3 days to induce *Cre*/*lox*P-mediated morphogenic gene excision. Embryos were then placed on maturation medium containing 50 mg/L of G418 and cultured at 28˚C in the dark for 10 days. Actively growing callus tissue was subcultured on fresh maturation medium at 28˚C in the dark for 14 days. About 33 days after infection, developing shoots were transferred to rooting medium containing 50 mg/L of G418 and incubated at 26˚C in a light chamber (16 h light/ 8 h dark) with 80 µmol/$m^2$/s for 10-14 days. Selected *in vitro* plantlets with developing roots (>2-3 cm) were shipped to Hawai'i, where they were transferred to sterile soilless mix (Sunshine Mix #4) in 10-cm pots and placed under a humi-dome with supplemental lighting (16 hr light, 8 hr dark) at room temperature (~26 ˚C) for about 10 days. After gradually removing the humi-dome and being fully exposed to air for 2-3 days, the acclimatized plants were transplanted into 7.5-L pots, moved to the University of Hawai'i greenhouse, and placed under a bench in the shade for another week before being moved to full light. Plants were grown under natural light (short-day conditions) and at maturity crossed as outlined in Figure S1.

3. Molecular analysis of edited plants

3.1 Sanger sequencing and identification of CRISPR edits. All sequence information corresponding to the target genes, *ZmCCT9* (Zm00042ab408960), *ZmCCT10* (Zm00042ab439330), and *ZmRap2.7* (Zm00042ab372190) were obtained from MaizeGDB genome assembly Zm-B73-REFERENCE-NAM-5.0. For molecular analysis of T0 transgenic plants, genomic DNA from leaf tissue was extracted using the EZNA Plant DNA DS Kit (Omega Bio-tek) according to the manufacturer's instructions. DNA quality and concentration were verified spectrophotometrically at 260/280 nm. To confirm T-DNA integration, PCR amplification was performed using primers specific to the selectable marker *neomycin phosphotransferase II* (*NPTII*) and *SpCas9* (Table S2). T-DNA copy number was determined using droplet digital PCR (QX200 Droplet Digital PCR system; Bio-Rad) using *Hin*dIII-digested genomic DNA as template per the manufacturer's instructions. Primers/probes specific to the transgenic selectable marker *NPTII*, and the endogenous reference gene MAPK kinase 1 (*ZmMEK1*; GenBank ZMU83625, Liu et al., 2011) were designed to detect the transgene copy number (Table S2). Target regions for *ZmCCT9*, *ZmCCT10*, and *ZmRap2.7* were amplified using gene-specific primers (Table S2). PCR products were purified using Exo-CIP$^{TM}$ Rapid PCR-Cleanup Kit using manufacturer's directions, sequenced by Sanger sequencing (Azenta/GeneWiz), and .ab1 trace files were analyzed by Tracking of Indels by Decomposition (TIDE v3.2.0; 50 bp window) and Inference of CRISPR Edits (ICE) to estimate indel frequencies (Brinkman et al. 2014; Conant et al. 2022).

3.2 PCR cloning and Sanger sequencing. To resolve specific allelic variants from T1 progeny, target gene edit regions were PCR amplified using gene-specific primers (Table S2) using T1 plant template DNA, amplicons were subcloned into the pGEM-T Easy vector (Promega), and up to 16 colonies per individual were sequenced by Sanger sequencing. Selected individuals in the T2 and T3 generations were screened by PCR followed by Sanger sequencing to confirm the allelic variants resolved in the T1 generation.

3.3 Cleaved Amplified Polymorphic Sequences (CAPS) assay (Matuszczak et al. 2020). CAPS assays were developed to screen for a complex 60 bp deletion with 27 bp insertion of novel sequence (hereafter called the 33 bp deletion) in *ZmCCT9* (*zmcct9*) and a 44 bp deletion in *ZmRap2.7* (*zmrap2.7*) using *Ava*II and *Asc*I restriction enzymes, respectively. PCR amplification products (Table S2) for *ZmCCT9* and *ZmRap2.7* were digested and analyzed by gel electrophoresis. Individuals homozygous for the deletion edits were identified by the presence of a single band, indicating no cleavage, wildtype individuals retained the restriction enzyme cleavage site, producing two bands, and heterozygous individuals produced three bands after digestion. Individuals in the T2 and T3 generation that lacked a wild-type digestion pattern were determined to be homozygous for the deletion allele and selected for subsequent analysis.

4. Phenotype analyses

4.1 Field phenotyping. The two fixed lines and Tzi8 inbred were grown as five replicates per genotype (1 row/replicate) with 20 kernels planted per row under standard maize cultivation conditions in Hawai'i (longest day and Iowa in summer 2025. Fields were planted on May 19 for Hawai'i and May 31 for Iowa. Day lengths in each location reached maximum on the summer solstice (June 21) which were 13.5 hrs. day/10.5 hrs. night in Hawai'i and 15.25 hrs. day/8.75 hrs. night in Iowa and declined gradually throughout the growing season. After maturity, several flowering traits were collected on all plants, which were numbered individually by row (n = 3 – 15 plants/rep) in both locations, including total leaf number (TLN), days to anthesis (DTA), and days to silking (DTS). In general, protocols were based on the Genomes2Fields standard operating procedures (SOP) manual (https://www.genomes2fields.org/resources/#sop).

4.2 Total leaf number. Measurement of total leaf number was done by marking the fifth leaf produced at the V4-V5 growth stage with an indelible marker. The mark was made near the leaf blade sheath junction, typically on the adaxial face of the leaf. At the V9-V10 growth stage, the 10th leaf was similarly marked, as leaf number 5 usually had senesced before the total number of leaves had been produced. After tassel emergence, total leaf number was calculated based on counting leaves from the uppermost marked leaf to the last leaf made which subtended the tassel (Supporting Data Tables S11 and S13).

4.3 Days to anthesis and silking. Daily observations were recorded during the flowering period for all plants in the field between 9 am to noon. The date of anthesis was recorded on individual plant tags if greater than 50% of the spikelets had anthers exserted from the central rachis (main spike) of the tassel. The date of silking was recorded on individual plant tags if 5-50 silk tips had exserted past the tips of the husk leaves on the uppermost ear (Supporting Data Tables S12 and S14).

4.4 Determining the floral transition. The developmental state of the shoot apical meristem (SAM) was determined by collecting whole V4 stage plants (16 days after planting) from random replicated rows from the Hawai'i nursery. The floral transition was determined by visual inspection after carefully dissecting mature and immature leaves from shoot apices to expose the SAM. Vegetative stage SAMs are proportionally domed shaped and initiate leaf primordia. Meristems that have elongated to a size approximately twice as tall as wide or have initiated visible branch meristems have passed the floral transition stage (Meng et al., 2011).

5 Data analyses

Traits were analyzed at the individual plant level for TLN, DTA, and DTS. Data analyses were performed in R (R Core Team, 2018) using linear mixed-effects models fit with the **lme4** and **lmerTest** packages. For each trait, the model (Y \sim \text{Genotype} \times \text{Location} + (1|\text{Location}:\text{Rep})) was fit, with Genotype, Location, and their interaction treated as fixed effects and replication (Rep) nested within Location included as a random effect. The overall significance of the fixed effects (Genotype, Location, and Genotype × Location) was evaluated using Type III F-tests from the mixed-effects model. Two biologically motivated contrasts were specified *a priori*: (i) Iowa versus Hawai'i within each genotype and (ii) pairwise genotype comparisons within each location. Estimated marginal means (EMMs) and their 95% confidence intervals were calculated using the **emmeans** package (v1.10.1), and pairwise comparisons were evaluated using t-tests based on the fitted mixed model. To control the false discovery rate associated with multiple comparisons, $P$-values were adjusted using the Benjamini-Hochberg false discovery rate (FDR) procedure, with statistical significance declared at an adjusted $P < 0.05$.

### Supplemental Figures

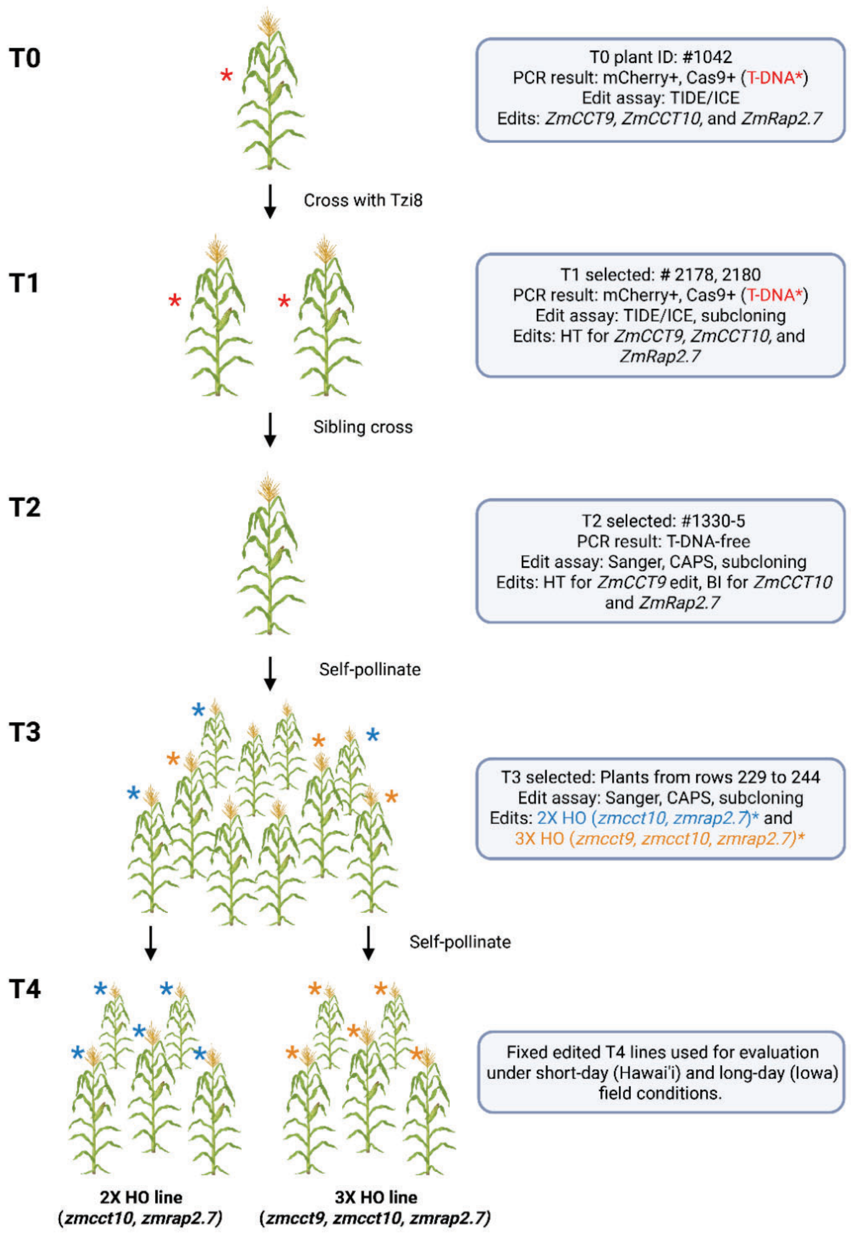

**Figure S1.** Crossing and genotyping overview from T0 to fixed T4 lines. Generation T0 to T4 (left) aligned with the crossing scheme (center) of selected genotypes that were determined by various molecular assays (right) to produce the fixed lines (blue, 2X HO and orange, 3X HO) used for field evaluations. Created in BioRender. Muszynski, M. (2026) https://BioRender.com/eyftsen

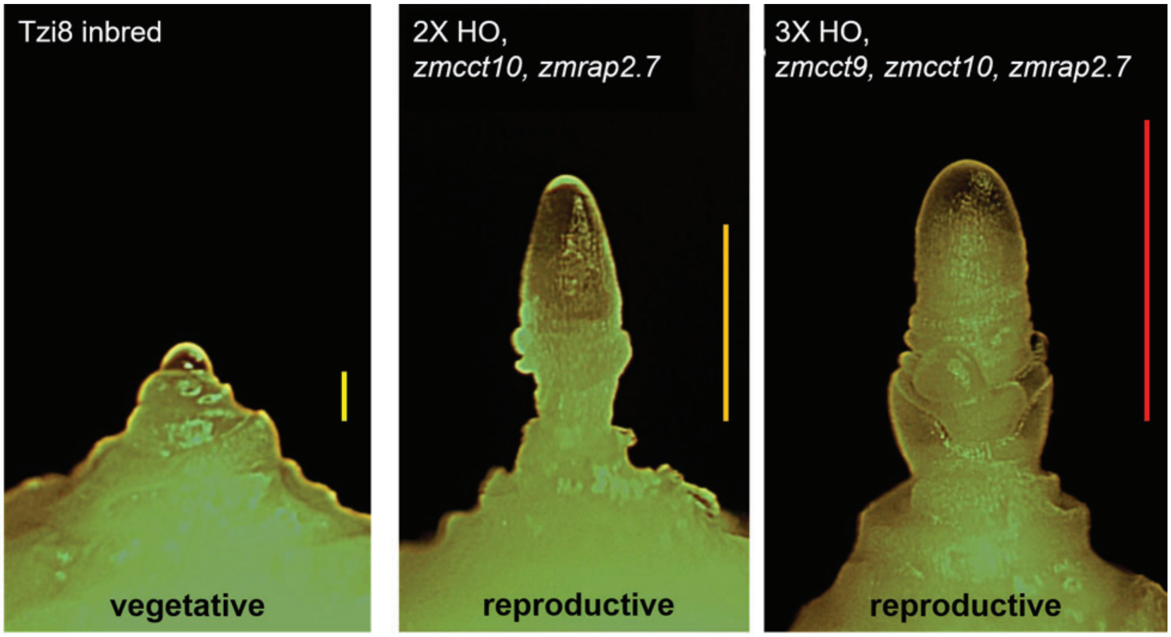

**Figure S2.** Developmental state of the shoot apical meristem (SAM) of Tzi8 and edited plants. Representative SAMs of Tzi8, 2X HO double-edited, and 3X HO triple-edited plants, collected 16 days after planting at growth stage V4 in short-days (Hawai'i) (left-to-right). The Tzi8 SAM is still in a vegetative state producing leaf primordia, while both edited plants have already transitioned to a reproductive state as their SAMs have become inflorescence meristems that have ceased making leaves and have initiated floral structures. Scale bars are 0.25 mm (yellow), 1.0 mm (orange), and 1.5 mm (red).

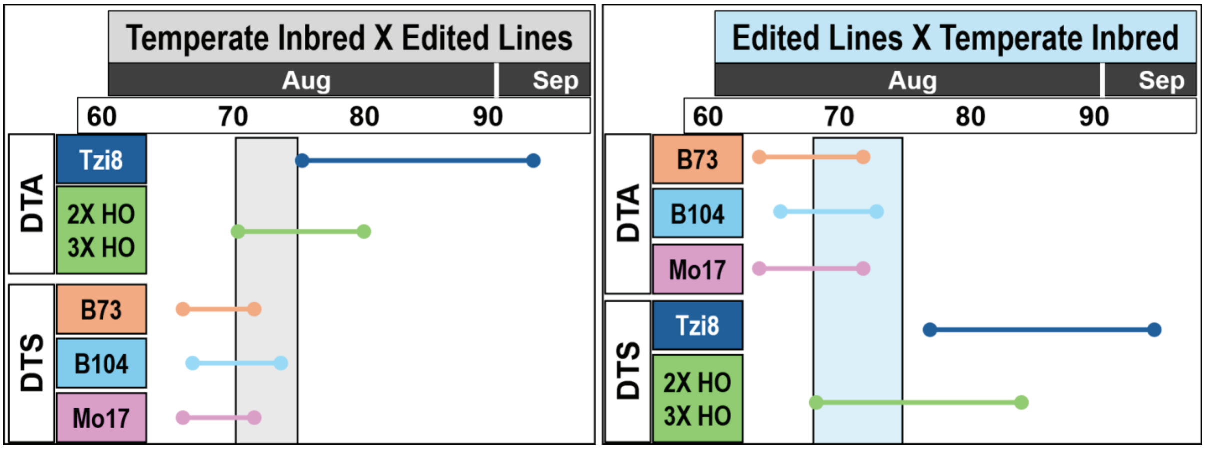

**Figure S3.** Time range for reciprocal cross pollination between tropical, gene edited tropical, and temperate lines under long-day conditions. (Top) The flowering intervals are aligned to the days after planting under typical long day (Iowa) summer field conditions. (Left) Days to anthesis (DTA) interval of Tzi8 and the two fixed edited lines (2X HO, 3X HO) aligned with days to silking (DTS) interval of three representative temperate inbreds (B73, B104, and Mo17). The gray rectangle indicates an overlap between the anthesis interval of the two fixed edited lines and the silking interval of temperate inbreds but not for Tzi8. (Right) DTS interval of Tzi8 and the two fixed edited lines (2X HO, 3X HO) aligned with the DTA interval of the three temperate inbreds. The blue rectangle indicates an overlap between the silking interval of the two fixed edited lines and the anthesis interval of the temperate inbreds but not for Tzi8. For tropical lines, the solid horizontal bars represent the observed flowering time ranges for DTA and DTS from this study. For temperate inbreds (B73, B104, and Mo17), the solid horizontal bars represent estimated marginal means (EMMs) for DTA and DTS calculated from previous Iowa summer field studies (Ghenov, 2026).

### Supplemental Tables

**Table S1. Summary of Tzi8 immature embryo transformation.**

| Experiment | #Emb | #Reg | #Non-excised | #Excised | #Escape | %TF |
|---|---|---|---|---|---|---|
| Exp 1 | 75 | 0 | 0 | **0** | 0 | 0 |
| Exp 2, Ear 1 | 64 | 1 | 0 | **0** | 1 | 0 |
| Exp 2, Ear 2 | 137 | 9 | 4 | **5** | 0 | 3.6 |
| Exp 2, Ear 3 | 136 | 0 | 0 | **0** | 0 | 0 |
| **Total** | **412** | **10** | **4** | **5** | **1** | **1.2** |

Emb, embryo; Reg, regenerated plantlets; TF, transformation frequency.

**Table S2. List of oligonucleotides used in this study.**

| Oligo name | Description | Sequence (5' - 3') |
|---|---|---|
| ZmCCT9-F1 | Forward PCR primer to amplify ZmCCT9 target region | CGAGCACATCATCATTCTC |
| ZmCCT9-R1 | Reverse PCR primer to amplify ZmCCT9 target region | CACAACCACGTAACACGAATC |
| ZmCCT9-SF1 | Forward primer for Sanger sequencing analysis of ZmCCT9 target region | CCGCGTATGTGGCGGCGTAG |
| ZmCCT10-F1 | Forward PCR primer to amplify ZmCCT10 target region | ATGGCAAGATCTCTTGGTACTTG |
| ZmCCT10-R1 | Reverse PCR primer to amplify ZmCCT10 target region | CCCAAGCTAGTCGATCCATC |
| ZmCCT10-SF1 | Forward Primer for Sanger sequencing analysis of ZmCCT10 target region | CGGGAGCAATACTTACGATGGTG |
| ZmRap2.7-F1 | Forward PCR primer to amplify ZmRap2.7 target region | GAAGTAGACACCACCAGTTCG |
| ZmRap2.7-R1 | Reverse PCR primer to amplify ZmRap2.7 target region | GTGGCTAGTTGTAGACAGTGG |
| ZmRap2.7-SF1 | Forward Primer for Sanger sequencing analysis of ZmRap2.7 target region | GGTGGCGAGGAGGAGGCGAT |
| ZmCCT9-gRNA1-F | Forward oligo for ZmCCT9 sgRNA1 | TGGCAGCCACATCCCGGGAAACCCG |
| ZmCCT9-gRNA1-R | Reverse oligo for ZmCCT9 sgRNA1 | AAACCGGGTTTCCCGGGATGTGGCT |
| ZmCCT10-gRNA1-F | Forward oligo for ZmCCT10 sgRNA1 | TGGCAGTCCCAAACGGGTGAAACGC |
| ZmCCT10-gRNA1-R | Reverse oligo for ZmCCT10 sgRNA1 | AAACGCGTTTCACCCGTTTGGGACT |
| ZmRap2.7-gRNA1-F | Forward oligo for ZmRap2.7 sgRNA1 | TGGCAGAGCTAGATCCCTCCTCCGC |
| ZmRap2.7-gRNA1-R | Reverse oligo for ZmRap2.7 sgRNA1 | AAACGCGGAGGAGGGATCTAGCTCT |
| MEK1 PrimeTime Primer 1 | Forward oligo for ZmMEK1 ddPCR assay | CTCAACTGCAGCCTCCCTAC |
| MEK1 PrimeTime Primer 2 | Reverse oligo for ZmMEK1 ddPCR assay | CACAGCCGATAAGACTGCAA |
| MEK1 PrimeTime Probe | MEK1 PrimeTime Probe (PrimeTime™ Assay Std Probe 5' HEX/ZEN™/3' IB™FQ) | /5HEX/AGAGGGAAC/ZEN/TATGCTG CATTCCGTGT/3IABkFQ/ |
| NPTII PrimeTime Primer 1 | Forward oligo for NPTII ddPCR assay | GATGGATTGCACGCAGGTTC |
| NPTII PrimeTime Primer 2 | Reverse oligo for NPTII ddPCR assay | GCCTCGTCCTGCAGTTCATT |
| NPTII PrimeTime Probe | NPTIIPrimeTime Probe for ddPCR | /56-FAM/CTCTGATGC/ZEN/CGCCG TG TTCC/3IABkFQ/ |

**Table S3. Summary of T0 gene edits detected by ICE analysis, T-DNA copy number, and T1 progeny seeds produced.**

| Plant ID | *ZmCCT9* | *ZmCCT10* | *ZmRap2.7* | T-DNA Copy# | T1 seeds |
|---|---|---|---|---|---|
| 1038 | -1 bp, 89.5% | +1 bp, 96.0% | n/a | 1 | Seed, OC |
| 1039 | -7 bp, 20.9%<br>-1 bp, 31.9% | +1 bp, 37.0% | +1 bp, 72.2% | 2 | Seed, OC |
| 1040 | -1 bp, 71.6% | +7 bp, 57.0% | -1 bp, 29.3%<br>+1 bp, 65.4% | 2 | No seed |
| 1041 | -18 bp, 19.2%<br>-12 bp, 26.6% | +7 bp, 49.6% | +1 bp, 42.1%<br>WT, 31.5% | 1 | No seed |
| **1042†** | -39 bp, 19.7%<br>-33 bp, 19.3% | +5 bp, 54.3% | +1 bp, 95.4% | 2 | Seed, OC |

Note: Percentage (%) next to the indel mutations indicates the knockout score reported by ICE analysis. OC, outcrossed to Tzi8. † T0 plant that produced the fixed lines evaluated in the field.

**Table S4. Summary of genotyping results of T1 progeny produced from the T0 event #1042**

| Plant ID | *ZmCCT9* | *ZmCCT10* | *ZmRap2.7* |
|---|---|---|---|
| 2167 | -33 bp (5/8) | -1 bp (4/6) | +1 bp (2/11) |
| 2176 | -60/+27 bp (12/17); CAPS | -1 bp (7/8) | +1 bp (5/11) |
| **2178†** | -1 bp (3/5) | -1 bp (10/21) | -44 bp (13/18); CAPS |
| **2180†** | -60/+27 bp (3/4); CAPS | +1 bp (8/10) | +1 bp (8/12) |

Note: Numbers in parentheses indicate the number of PCR amplicon clones sequenced. CAPS, Cleaved Amplified Polymorphic Sequences assay using the *Ava*II restriction enzyme for the -60/+27 bp complex *ZmCCT9* edit and *Acs*I restriction enzyme for the -44 bp *ZmRap2.7* deletion alleles, respectively. †T1 plants sib-pollinated that produced the fixed lines evaluated in the field

**Table S5. Summary of T1 gene editing alleles identified by subclone sequencing**

| T1 ID | Gene | DNA sequence (5′→3′) | Allele/edit |
|---|---|---|---|
| **2167** | ***ZmCCT9*** | ACCTCAACCGCGGGTTTCCCGGGATGTGGCACCAGCACCACCAGCAGCGC | **WT** |
| | | ACCTCAACCGC**GG**GGGTTTCCCGGGATGTGGCACCAGCACCACCAGCAGCGC | +2 bp |
|  |  | ACCTCAACCG---------------------------------GCAGC | −33 bp |
| | ***ZmCCT10*** | GACCAGCCTGCGTTTCACCCGTTTGGGACACCACAGT | **WT** |
| | | GACCAGCCTGC-TTTCACCCGTTTGGGACACCACAGT | −1 bp |
|  |  | GACCAGCCTGCGTT----------GGGACACCACAGT | −10 bp |
| | ***ZmRap2.7*** | GCCCGCGCCGGCGGAGGAGGGATCTAGCTCAACGCC | **WT** |
| | | GCCCGCGCCGGCG**T**GAGGAGGGATCTAGCTCAACGCC | +1 bp |
| | | GCCCGCGCCGGCG**A**GAGGAGGGATCTAGCTCAACGCC | +1 bp |
| | | GCCCGCGCCGG---GAGGAGGGATCTAGCTCAACGCC | −3 bp |
| | | GCCCGCGCCGGC--GAGGAGGGATCTAGCTCAACGCC | −2 bp |
|  |  | GCCCGCGCCGGC--------GGATCTAGCTCAACGCC | −8 bp |
| | | GCCCGCGCCGGCG-GAGGAGGGATCTAGCTCAACGCC | −1 bp |
| **2176** | ***ZmCCT9*** | ACCTCAACCGCGGGTTTCCCGGGATGTGGCACCAGCACCACCAGCAGCGC | **WT** |
|  |  | ----------------------CCGGGATGTGGCACCAGCACCACCAGCAGC | −22 bp |
| | | **GGGGATACAAATTTGATCCCGGGATGT**--//--GGATGTGGCACCAG | -60/+27 bp / |
| | ***ZmCCT10*** | GACCAGCCTGCGTTTCACCCGTTTGGGACACCACAGT | **WT** |
| | | GACCAGCCTGC-TTTCACCCGTTTGGGACACCACAGT | −1 bp |
| | | GACCAGCCTGC**A-**TTCACCCGTTTGGGACACCACAGT | sub/−1 bp |
| | ***ZmRap2.7*** | GCCCGCGCCGGCGGAGGAGGGATCTAGCTCAACGCC | **WT** |
| | | GCCCGCGCCGGCG**C**GAGGAGGGATCTAGCTCAACGCC | +1 bp |
| | | GCCCGCGCCGGCG**A**GAGGAGGGATCTAGCTCAACGCC | +1 bp |
|  |  | GGG----------------//----------------CC | −44 bp |
| | | GCCCGCGCCGGCG--AGGAGGGATCTAGCTCAACGCC | −2 bp |
|  |  | GCCCGCGCCG-----AGGAGGGATCTAGCTCAACGCC | −5 bp |
| **2178†** | ***ZmCCT9*** | ACCTCAACCGCGGGTTTCCCGGGATGTGGCACCAGCACCACCAGCAGCGC | **WT** |
|  |  | ACCTCAACCG---------------------------------GCAGC | −33 bp |
| | | ACCTCAACCGCG-GTTTCCCGGGATGTGGCACCAGCNCCACCAGCAGC | −1 bp |
| | ***ZmCCT10*** | GACCAGCCTGCGTTTCACCCGTTTGGGACACCACAGT | **WT** |
| | | GACCAGCCTGC-TTTCACCCGTTTGGGACACCACAGT | −1 bp‡ |
|  |  | GACCA------------CCCGTTTGGGACACCACAGT | −12 bp |
|  |  | GACC---------------CGTTTGGGACACCACAGT | −15 bp |
| | ***ZmRap2.7*** | GCCCGCGCCGGCGGAGGAGGGATCTAGCTCAACGCC | **WT** |
|  |  | GGG----------------//----------------CC | −44 bp |
| | | GCCCGCGCCGGCG--AGGAGGGATCTAGCTCAACGCC | −2 bp |
| **2180†** | ***ZmCCT9*** | ACCTCAACCGCGGGTTTCCCGGGATGTGGCACCAGCACCACCAGCAGCGC | **WT** |
| | | **GGGGATACAAATTTGATCCCGGGATGT**--//--GGATGTGGCACCAG | -60/+27 bp‡ |
|  |  | ----//------------GGTTTCCCGGGATGTGGCACCGCACCACCAGCAGC | −44 bp |
| | ***ZmCCT10*** | GACCAGCCTGCGTTTCACCCGTTTGGGACACCACAGT | **WT** |
| | | GACCAGCCTGCG**A**TTTCACCCGTTTGGGACACCACAGT | +1 bp |
| | | GACCAGCCTGCG**AC**TTCACCCGTTTGGGACACCACAGT | sub/+1 bp |
| | | GACCAGCCTGCG**T**TTTCACCCGTTTGGGACACCACAGT | +1 bp‡ |
| | ***ZmRap2.7*** | GCCCGCGCCGGCGGAGGAGGGATCTAGCTCAACGCC | **WT** |

**Table S5. Summary of T1 gene editing alleles identified by subclone sequencing**

| T1 ID | Gene | DNA sequence (5′→3′) | Allele/edit |
|---|---|---|---|
| | | GCCCGCGCCGGCG**C**GAGGAGGGATCTAGCTCAACGCC | +1 bp |
| | | GCCCGCGCCGGCG**T**GAGGAGGGATCTAGCTCAACGCC | +1 bp |
| | | GCCCGCGC**T**GGCG**C**GAGGAGGGATCTAGCTCAACGCC | sub/+1 bp |
|  |  | CGGC--------------------//----------------CGACCT | −256 bp |

Note: Wild type (WT) alleles are bolded, PAM sequences are underlined, and insertions, substitutions (sub), and complex rearrangements are bolded with yellow highlights. †T1 plants sib-pollinated that produced the fixed lines evaluated in the field. ‡ Alleles identified as homozygous or biallelic in the fixed lines.

**Table S6. Summary of genotyping results of T-DNA free T2 and T3 plants.**

| T1 ID† | T2 ID‡ | T3 IDs§ | *ZmCCT9* | | *ZmCCT10* | | *ZmRap2.7* | |
|---|---|---|---|---|---|---|---|---|
| | | | **T2 genotype** | **T3 genotype** | **T2 genotype** | **T3 genotype** | **T2 genotype** | **T3 genotype** |
| 2176/ Tzi8 | 1324-8 | 196 to 214 | -22 bp; HT | -22 bp; HO segregating | -1 bp; HT | -1 bp; HT | +1 bp; HT | +1 bp; HT or HO |
| 2178/ Tzi8 | 1328-12 | 181 to 195 | -1 bp; HT | -1 bp; HO segregating | -1 bp; HT | -1 bp; HT | -44 bp; HT | -44 bp; HT or HO |
| 2180/ 2178 | 1330-1 | 215 to 228 | -33 bp; HT | -60/+27 bp; HO segregating | +1 bp; HO fixed | +1 bp; HO fixed | +1 bp (C)/+1 bp (A); BI | +1 bp (C)/+1 bp (A); HO or BI |
| 2180/ 2178 | **1330-5**¶ | 229 to 244 | -33 bp; HT | -60/+27 bp; HO segregating | +1 bp/-1 bp; BI | +1 bp/-1 bp; HO or BI | +1 bp (C)/+1 bp (T); BI | +1 bp (C)/+1 bp (T); HO or BI |

Note: HT, Heterozygous; HO, Homozygous; BI, Biallelic. † T1 plant ID and cross to Tzi8 or sib pollination. ‡ T-DNA free T2 plants that were self-pollinated. § These IDs are row numbers with 8 - 13 T3 plants each. ¶ T-DNA free T2 plant that produced the fixed lines evaluated in the field.

**Table S7. Total leaf number (TLN) estimated marginal means (EMM) and statitical analysis used to produce the violin plots in Figure 1F.**

| Location | Genotype | EMM | SE | 95% CI (L) | 95% CI (U) | n | Letters |
|---|---|---|---|---|---|---|---|
| Hawaii | Tzi8 | 19.26 | 0.21 | 18.85 | 19.67 | 36 | b |
| Hawaii | 2X HO | 15.72 | 0.18 | 15.35 | 16.10 | 48 | a |
| Hawaii | 3X HO | 15.68 | 0.16 | 15.34 | 16.02 | 69 | a |
| Iowa | Tzi8 | 22.91 | 0.19 | 22.52 | 23.30 | 42 | c |
| Iowa | 2X HO | 20.13 | 0.16 | 19.79 | 20.47 | 67 | b |
| Iowa | 3X HO | 19.41 | 0.16 | 19.07 | 19.75 | 71 | a |

2X HO is the fixed double-edited *zmcct10, zmrap2.7* line.

3X HO is the fixed triple-edited *zmcct9, zmcct10, zmrap2.7* line.

**Table S8. Total leaf number (TLN) estimated marginal mean differences between genotypes by location.**

| Location | Genotype A | Genotype B | A - B (diff) | SE (diff) | df (diff) | t (diff) | 95% CI diff | 95% CI diff | p adj (BH) | sig |
|---|---|---|---|---|---|---|---|---|---|---|
| Hawaii | Tzi8 | 3X HO | 3.58 | 0.22 | 322.19 | 16.18 | 3.05 | 4.11 | 4.84E-43 | **** |
| Hawaii | Tzi8 | 2X HO | 3.54 | 0.24 | 326.28 | 14.77 | 2.96 | 4.11 | 5.49E-38 | **** |
| Hawaii | 2X HO | 3X HO | 0.04 | 0.20 | 323.11 | 0.20 | -0.45 | 0.53 | 0.8396779 | ns |
| Iowa | Tzi8 | 3X HO | 3.50 | 0.21 | 321.06 | 16.75 | 3.00 | 4.00 | 3.10E-45 | **** |
| Iowa | Tzi8 | 2X HO | 2.78 | 0.21 | 323.51 | 13.11 | 2.27 | 3.29 | 1.36E-31 | **** |
| Iowa | 2X HO | 3X HO | 0.72 | 0.18 | 320.31 | 3.94 | 0.28 | 1.16 | 1.01E-04 | *** |

2X HO is the fixed double-edited *zmcct10, zmrap2.7* line.

3X HO is the fixed triple-edited *zmcct9, zmcct10, zmrap2.7* line.

**Table S9. Flowering traits (DTA, DTS) estimated marginal means (EMM) and statitical analysis used to produce the violin plots in Figure 1F.**

| Trait | Location | Genotype | EMM | SE | 95% CI (L) | 95% CI (U) | n | Letters |
|---|---|---|---|---|---|---|---|---|
| DTA | Hawaii | Tzi8 | 57.26 | 0.76 | 55.68 | 58.83 | 28 | b |
| DTA | Hawaii | 2X HO | 53.21 | 0.70 | 51.73 | 54.70 | 41 | a |
| DTA | Hawaii | 3X HO | 52.30 | 0.65 | 50.89 | 53.72 | 62 | a |
| DTA | Iowa | Tzi8 | 82.64 | 0.74 | 81.10 | 84.19 | 31 | b |
| DTA | Iowa | 2X HO | 74.20 | 0.65 | 72.79 | 75.62 | 62 | a |
| DTA | Iowa | 3X HO | 74.57 | 0.64 | 73.16 | 75.97 | 69 | a |
| GDD_DTA | Hawaii | Tzi8 | 1696.88 | 20.40 | 1654.35 | 1739.41 | 28 | b |
| GDD_DTA | Hawaii | 2X HO | 1575.33 | 18.87 | 1535.07 | 1615.59 | 41 | a |
| GDD_DTA | Hawaii | 3X HO | 1548.31 | 17.59 | 1509.72 | 1586.91 | 62 | a |
| GDD_DTA | Iowa | Tzi8 | 2000.19 | 19.90 | 1958.44 | 2041.95 | 31 | b |
| GDD_DTA | Iowa | 2X HO | 1790.62 | 17.63 | 1751.97 | 1829.27 | 62 | a |
| GDD_DTA | Iowa | 3X HO | 1801.27 | 17.34 | 1762.96 | 1839.58 | 69 | a |
| DTS | Hawaii | Tzi8 | 56.38 | 0.63 | 55.09 | 57.67 | 33 | b |
| DTS | Hawaii | 2X HO | 54.09 | 0.56 | 52.92 | 55.26 | 52 | a |
| DTS | Hawaii | 3X HO | 53.28 | 0.53 | 52.15 | 54.40 | 64 | a |
| DTS | Iowa | Tzi8 | 81.88 | 0.53 | 80.76 | 83.01 | 65 | b |
| DTS | Iowa | 2X HO | 74.70 | 0.53 | 73.57 | 75.83 | 62 | a |
| DTS | Iowa | 3X HO | 74.51 | 0.52 | 73.39 | 75.63 | 67 | a |
| GDD_DTS | Hawaii | Tzi8 | 1669.45 | 17.35 | 1633.65 | 1705.25 | 33 | b |
| GDD_DTS | Hawaii | 2X HO | 1601.35 | 15.61 | 1568.24 | 1634.45 | 52 | a |
| GDD_DTS | Hawaii | 3X HO | 1577.42 | 14.88 | 1545.36 | 1609.47 | 64 | a |
| GDD_DTS | Iowa | Tzi8 | 1985.39 | 14.83 | 1953.41 | 2017.37 | 65 | b |
| GDD_DTS | Iowa | 2X HO | 1803.12 | 14.96 | 1770.95 | 1835.28 | 62 | a |
| GDD_DTS | Iowa | 3X HO | 1799.29 | 14.77 | 1767.37 | 1831.21 | 67 | a |

2X HO is the fixed double-edited *zmcct10, zmrap2.7* line.

3X HO is the fixed triple-edited *zmcct9, zmcct10, zmrap2.7* line.

**Table S10. Flowering traits (DTA, DTS) estimated marginal mean differences between genotypes by location.**

| Trait | Location | Genotype A | Genotype B | A - B (diff) | SE (diff) | df (diff) | t (diff) | 95% CI diff | 95% CI diff | p adj (BH) | sig |
|---|---|---|---|---|---|---|---|---|---|---|---|
| DTA | Hawaii | Tzi8 | 3X HO | 4.95 | 0.64 | 279.82 | 7.74 | 3.41 | 6.50 | 5.53E-13 | **** |
| DTA | Hawaii | Tzi8 | 2X HO | 4.05 | 0.70 | 281.58 | 5.81 | 2.37 | 5.72 | 2.51E-08 | **** |
| DTA | Hawaii | 2X HO | 3X HO | 0.91 | 0.57 | 280.28 | 1.60 | -0.46 | 2.28 | 0.110106841 | ns |
| DTA | Iowa | Tzi8 | 2X HO | 8.44 | 0.62 | 280.78 | 13.55 | 6.94 | 9.94 | 4.87E-32 | **** |
| DTA | Iowa | Tzi8 | 3X HO | 8.08 | 0.61 | 279.78 | 13.27 | 6.61 | 9.54 | 2.40E-31 | **** |
| DTA | Iowa | 2X HO | 3X HO | -0.36 | 0.49 | 280.36 | -0.73 | -1.55 | 0.83 | 0.463715745 | ns |
| GDD_DTA | Hawaii | Tzi8 | 3X HO | 148.57 | 16.67 | 279.74 | 8.91 | 108.41 | 188.73 | 2.01E-16 | **** |
| GDD_DTA | Hawaii | Tzi8 | 2X HO | 121.55 | 18.13 | 281.36 | 6.70 | 77.89 | 165.22 | 1.65E-10 | **** |
| GDD_DTA | Hawaii | 2X HO | 3X HO | 27.02 | 14.77 | 280.17 | 1.83 | -8.57 | 62.60 | 0.068511399 | ns |
| GDD_DTA | Iowa | Tzi8 | 2X HO | 209.57 | 16.22 | 280.62 | 12.92 | 170.50 | 248.64 | 8.45E-30 | **** |
| GDD_DTA | Iowa | Tzi8 | 3X HO | 198.93 | 15.84 | 279.70 | 12.56 | 160.77 | 237.08 | 8.36E-29 | **** |
| GDD_DTA | Iowa | 2X HO | 3X HO | -10.64 | 12.87 | 280.23 | -0.83 | -41.63 | 20.34 | 0.408725662 | ns |
| DTS | Hawaii | Tzi8 | 3X HO | 3.11 | 0.60 | 331.70 | 5.15 | 1.66 | 4.56 | 1.33E-06 | **** |
| DTS | Hawaii | Tzi8 | 2X HO | 2.29 | 0.63 | 334.60 | 3.62 | 0.77 | 3.82 | 5.06E-04 | *** |
| DTS | Hawaii | 2X HO | 3X HO | 0.81 | 0.53 | 333.59 | 1.54 | -0.46 | 2.09 | 0.124473508 | ns |
| DTS | Iowa | Tzi8 | 3X HO | 7.37 | 0.49 | 329.95 | 15.13 | 6.20 | 8.55 | 3.71E-39 | **** |
| DTS | Iowa | Tzi8 | 2X HO | 7.18 | 0.50 | 329.61 | 14.47 | 5.99 | 8.38 | 6.61E-37 | **** |
| DTS | Iowa | 2X HO | 3X HO | 0.19 | 0.50 | 331.10 | 0.38 | -1.00 | 1.38 | 0.702250948 | ns |
| GDD_DTS | Hawaii | Tzi8 | 3X HO | 92.03 | 15.57 | 331.20 | 5.91 | 54.56 | 129.50 | 2.55E-08 | **** |
| GDD_DTS | Hawaii | Tzi8 | 2X HO | 68.10 | 16.35 | 333.77 | 4.17 | 28.76 | 107.44 | 5.95E-05 | **** |
| GDD_DTS | Hawaii | 2X HO | 3X HO | 23.93 | 13.65 | 332.86 | 1.75 | -8.92 | 56.78 | 0.080602228 | ns |
| GDD_DTS | Iowa | Tzi8 | 3X HO | 186.10 | 12.59 | 329.75 | 14.79 | 155.81 | 216.39 | 7.96E-38 | **** |
| GDD_DTS | Iowa | Tzi8 | 2X HO | 182.27 | 12.82 | 329.48 | 14.22 | 151.43 | 213.12 | 6.19E-36 | **** |
| GDD_DTS | Iowa | 2X HO | 3X HO | 3.83 | 12.79 | 330.67 | 0.30 | -26.96 | 34.61 | 0.765093912 | ns |

2X HO is the fixed double-edited *zmcct10, zmrap2.7* line.

3X HO is the fixed triple-edited *zmcct9, zmcct10, zmrap2.7* line.

**Table S11. Total leaf number from Hawai'i summer 2025 field.**

| Rep | GENOTYPE | P#1 | P#2 | P#3 | P#4 | P#5 | P#6 | P#7 | P#8 | P#9 | P#10 | P#11 | P#12 | P#13 | P#14 | P#15 | P#16 |
|---|---|---|---|---|---|---|---|---|---|---|---|---|---|---|---|---|---|
| **1** | 2X HO | 15 | 16 | 15 | 16 | 16 | - | - | - | 15 | - | 15 | 15 | 16 | No mark | 16 | No tassel |
| **2** | 2X HO | 16 | 15 | 15 | 15 | 17 | 16 | 15 | 17 | 15 | 16 | 18 | 16 | - | - | - | - |
| **3** | 2X HO | 15 | 16 | 17 | 17 | 16 | - | - | - | - | - | - | - | - | - | - | - |
| **4** | 2X HO | 16 | 15 | 16 | 15 | 15 | 16 | 16 | 17 | 16 | 15 | 16 | 17 | - | - | - | - |
| **5** | 2X HO | 15 | 15 | 15 | 16 | 17 | 14 | 15 | 17 | 15 | - | - | - | - | - | - | - |
| **1** | 3X HO | 16 | 17 | 16 | 16 | 18 | 17 | 17 | 15 | 15 | 14 | 16 | 15 | 15 | 15 | - | - |
| **2** | 3X HO | 16 | 16 | 15 | 15 | 15 | 16 | 16 | 16 | 16 | 15 | 15 | 19 | 16 | 15 | 16 | - |
| **3** | 3X HO | 17 | 14 | 17 | 16 | 16 | 17 | 16 | 15 | 15 | 15 | 16 | 15 | 16 | - | 14 | - |
| **4** | 3X HO | 15 | 16 | 17 | 16 | 15 | 15 | 15 | 16 | 16 | 15 | 16 | 17 | - | - | - | - |
| **5** | 3X HO | 17 | 15 | 17 | 14 | 16 | 13 | 16 | 15 | 15 | 15 | 15 | 16 | 16 | 15 | - | - |
| **1** | Tzi8 | - | 21 | 20 | 19 | 19 | 20 | 19 | - | - | - | - | - | - | - | - | - |
| **2** | Tzi8 | 21 | 19 | 20 | - | 20 | 21 | 19 | - | 20 | 21 | 20 | 20 | - | - | - | - |
| **3** | Tzi8 | 20 | 14 | 19 | 14 | 20 | 20 | 20 | 20 | 20 | - | - | - | - | - | - | - |
| **4** | Tzi8 | 20 | 21 | 20 | - | - | - | - | - | - | - | - | - | - | - | - | - |
| **5** | Tzi8 | 20 | 17 | 19 | 20 | 14 | 19 | - | 19 | 18 | - | - | - | - | - | - | - |

2X HO is the fixed double-edited *zmcct10, zmrap2.7* line.

3X HO is the fixed triple-edited *zmcct9, zmcct10, zmrap2.7* line.

**Table S12. Flowering traits from Hawai'i summer 2025 field.**

| Rep | GENOTYPE | Trait | P#1 | P#2 | P#3 | P#4 | P#5 | P#6 | P#7 | P#8 | P#9 | P#10 | P#11 | P#12 | P#13 | P#14 | P#15 | P#16 |
|---|---|---|---|---|---|---|---|---|---|---|---|---|---|---|---|---|---|---|
| 1 | 2X HO | DTA | - | 52 | | 57 | 53 | 54 | - | - | - | - | 55 | | | 56 | 55 | - |
| 2 | 2X HO | DTA | - | 53 | 54 | 56 | 57 | 57 | 57 | 56 | - | - | - | 53 | | | | |
| 3 | 2X HO | DTA | 54 | 51 | 58 | 55 | 51 | | | | | | | | | | | |
| 4 | 2X HO | DTA | 52 | 49 | 51 | 49 | 51 | 53 | 52 | 53 | 49 | 49 | 52 | 52 | | | | |
| 5 | 2X HO | DTA | 53 | 53 | 50 | 50 | 52 | 53 | 51 | 54 | 52 | | | | | | | |
| 1 | 3X HO | DTA | 53 | - | - | 61 | 53 | 51 | 56 | 51 | 52 | 49 | 51 | 51 | - | 50 | | |
| 2 | 3X HO | DTA | 56 | 54 | 53 | 53 | 53 | 53 | 53 | 52 | 54 | 2/21 | 49 | - | 59 | 52 | 53 | |
| 3 | 3X HO | DTA | 54 | 53 | - | 53 | 55 | 57 | 53 | 53 | - | 51 | 53 | - | 53 | - | 52 | |
| 4 | 3X HO | DTA | 49 | 49 | 55 | 49 | 50 | 49 | 49 | 50 | 53 | 50 | 52 | 57 | | | | |
| 5 | 3X HO | DTA | 51 | 49 | 54 | 54 | 52 | 49 | 54 | 51 | 49 | 51 | 49 | 51 | 53 | 53 | | |
| 1 | Tzi8 | DTA | - | 56 | 56 | 56 | 57 | 58 | 55 | | | | | | | | | |
| 2 | Tzi8 | DTA | | 55 | 58 | - | 61 | 63 | 57 | | 59 | 59 | | 60 | | | | |
| 3 | Tzi8 | DTA | - | | 57 | - | 57 | 57 | 58 | - | 57 | | | | | | | |
| 4 | Tzi8 | DTA | 57 | 58 | 58 | | | | | | | | | | | | | |
| 5 | Tzi8 | DTA | 56 | - | 58 | 58 | - | 57 | - | 56 | 55 | | | | | | | |
| 1 | 2X HO | DTS | 54 | 53 | 56 | 51 | 52 | 54 | 53 | 52 | 53 | 53 | 52 | 58 | 54 | 53 | 53 | 53 |
| 2 | 2X HO | DTS | 56 | 58 | 56 | 56 | 56 | 56 | 57 | 59 | 56 | 56 | - | 55 | | | | |
| 3 | 2X HO | DTS | 55 | 51 | 59 | 56 | 52 | | | | | | | | | | | |
| 4 | 2X HO | DTS | 52 | 52 | 54 | 52 | 52 | 55 | 54 | 54 | 50 | 51 | 52 | 54 | | | | |
| 5 | 2X HO | DTS | 54 | 54 | 50 | 51 | 53 | 57 | 53 | 55 | - | | | | | | | |
| 1 | 3X HO | DTS | 54 | 52 | 52 | 52 | 54 | 53 | 55 | 50 | 52 | 50 | 50 | 51 | 57 | 52 | | |
| 2 | 3X HO | DTS | - | 57 | 56 | - | - | 55 | 54 | 56 | 58 | 52 | 52 | 64 | 52 | - | 54 | |
| 3 | 3X HO | DTS | 56 | 53 | 60 | 54 | 50 | 58 | 57 | 54 | - | 51 | 51 | 51 | 53 | 54 | 52 | |
| 4 | 3X HO | DTS | 49 | 49 | - | 50 | 51 | 50 | 50 | 51 | 53 | 50 | 52 | 57 | | | | |
| 5 | 3X HO | DTS | 52 | 51 | 55 | 55 | 54 | 53 | 56 | 52 | 51 | 51 | 58 | 54 | 54 | 53 | | |
| 1 | Tzi8 | DTS | - | 56 | 56 | 56 | 55 | 58 | 56 | | | | | | | | | |
| 2 | Tzi8 | DTS | 57 | 55 | 59 | - | 57 | 59 | 56 | | 58 | 58 | 56 | 57 | | | | |
| 3 | Tzi8 | DTS | - | 57 | 57 | - | 56 | 56 | 57 | 55 | 56 | | | | | | | |
| 4 | Tzi8 | DTS | 55 | 57 | 56 | | | | | | | | | | | | | |
| 5 | Tzi8 | DTS | 57 | - | 58 | 58 | 59 | 57 | - | 56 | 54 | | | | | | | |
| 1 | 2X HO | GDD_DTA | - | 1539 | | 1689 | 1568 | 1597 | - | - | - | - | 1628.5 | | | 1657.5 | 1628.5 | - |
| 2 | 2X HO | GDD_DTA | - | 1568 | 1597 | 1657.5 | 1689 | 1689 | 1689 | 1657.5 | - | - | - | 1568 | | | | |
| 3 | 2X HO | GDD_DTA | 1597 | 1510.5 | 1719.5 | 1628.5 | 1510.5 | | | | | | | | | | | |
| 4 | 2X HO | GDD_DTA | 1539 | 1451 | 1510.5 | 1451 | 1510.5 | 1568 | 1539 | 1568 | 1451 | 1451 | 1539 | 1539 | | | | |
| 5 | 2X HO | GDD_DTA | 1568 | 1568 | 1481 | 1481 | 1539 | 1568 | 1510.5 | 1597 | 1539 | | | | | | | |
| 1 | 3X HO | GDD_DTA | 1568 | - | - | 1811.5 | 1568 | 1510.5 | 1657.5 | 1510.5 | 1539 | 1451 | 1510.5 | 1510.5 | - | 1481 | | |
| 2 | 3X HO | GDD_DTA | 1657.5 | 1597 | 1568 | 1568 | 1568 | 1568 | 1568 | 1539 | 1597 | 1539 | 1451 | - | 1750 | 1539 | 1568 | |
| 3 | 3X HO | GDD_DTA | 1597 | 1568 | - | 1568 | 1628.5 | 1689 | 1568 | 1568 | - | 1510.5 | 1568 | - | 1568 | - | 1539 | |
| 4 | 3X HO | GDD_DTA | 1451 | 1451 | 1628.5 | 1451 | 1481 | 1451 | 1451 | 1481 | 1568 | 1481 | 1539 | 1689 | | | | |
| 5 | 3X HO | GDD_DTA | 1510.5 | 1451 | 1597 | 1597 | 1539 | 1451 | 1597 | 1510.5 | 1451 | 1510.5 | 1451 | 1510.5 | 1568 | 1568 | | |
| 1 | Tzi8 | GDD_DTA | - | 1657.5 | 1657.5 | 1657.5 | 1689 | 1719.5 | 1628.5 | | | | | | | | | |
| 2 | Tzi8 | GDD_DTA | | 1628.5 | 1719.5 | - | 1811.5 | 1874 | 1689 | | 1750 | 1750 | | 1780.5 | | | | |
| 3 | Tzi8 | GDD_DTA | - | | 1689 | - | 1689 | 1689 | 1719.5 | - | 1689 | | | | | | | |
| 4 | Tzi8 | GDD_DTA | 1689 | 1719.5 | 1719.5 | | | | | | | | | | | | | |
| 5 | Tzi8 | GDD_DTA | 1657.5 | - | 1719.5 | 1719.5 | | 1689 | - | 1657.5 | 1628.5 | | | | | | | |
| 1 | 2X HO | GDD_DTS | 1597 | 1568 | 1657.5 | 1510.5 | 1539 | 1597 | 1568 | 1539 | 1568 | 1568 | 1539 | 1719.5 | 1597 | 1568 | 1568 | 1568 |
| 2 | 2X HO | GDD_DTS | 1657.5 | 1719.5 | 1657.5 | 1657.5 | 1657.5 | 1657.5 | 1689 | 1750 | 1657.5 | 1657.5 | - | 1628.5 | | | | |
| 3 | 2X HO | GDD_DTS | 1628.5 | 1510.5 | 1750 | 1657.5 | 1539 | | | | | | | | | | | |

| | | | | | | | | | | | | | | | | | |
|---|---|---|---|---|---|---|---|---|---|---|---|---|---|---|---|---|---|
| **4** | 2X HO | GDD_DTS | 1539 | 1539 | 1597 | 1539 | 1539 | 1628.5 | 1597 | 1597 | 1481 | 1510.5 | 1539 | 1597 | | | |
| **5** | 2X HO | GDD_DTS | 1597 | 1597 | 1481 | 1510.5 | 1568 | 1689 | 1568 | 1628.5 | - | | | | | | |
| **1** | 3X HO | GDD_DTS | 1597 | 1539 | 1539 | 1539 | 1597 | 1568 | 1628.5 | 1481 | 1539 | 1481 | 1481 | 1510.5 | 1689 | 1539 | |
| **2** | 3X HO | GDD_DTS | - | 1689 | 1657.5 | - | - | 1628.5 | 1597 | 1657.5 | 1719.5 | 1539 | 1539 | 1905 | 1539 | - | 1597 |
| **3** | 3X HO | GDD_DTS | 1657.5 | 1568 | 1780.5 | 1597 | 1481 | 1719.5 | 1689 | 1597 | - | 1510.5 | 1510.5 | 1510.5 | 1568 | 1597 | 1539 |
| **4** | 3X HO | GDD_DTS | 1451 | 1451 | - | 1481 | 1510.5 | 1481 | 1481 | 1510.5 | 1568 | 1481 | 1539 | 1689 | | | |
| **5** | 3X HO | GDD_DTS | 1539 | 1510.5 | 1628.5 | 1628.5 | 1597 | 1568 | 1657.5 | 1539 | 1510.5 | 1510.5 | 1719.5 | 1597 | 1597 | 1568 | |
| **1** | Tzi8 | GDD_DTS | - | 1657.5 | 1657.5 | 1657.5 | 1628.5 | 1719.5 | 1657.5 | | | | | | | | |
| **2** | Tzi8 | GDD_DTS | 1689 | 1628.5 | 1750 | - | 1689 | 1750 | 1657.5 | | 1719.5 | 1719.5 | 1657.5 | 1689 | | | |
| **3** | Tzi8 | GDD_DTS | - | 1689 | 1689 | - | 1657.5 | 1657.5 | 1689 | 1628.5 | 1657.5 | | | | | | |
| **4** | Tzi8 | GDD_DTS | 1628.5 | 1689 | 1657.5 | | | | | | | | | | | | |
| **5** | Tzi8 | GDD_DTS | 1689 | - | 1719.5 | 1719.5 | 1750 | 1689 | - | 1657.5 | 1597 | | | | | | |

2X HO is the fixed double-edited *zmcct10, zmrap2.7* line.

3X HO is the fixed triple-edited *zmcct9, zmcct10, zmrap2.7* line.

DTA, days to anthesis; DTS, days to silking, GDD, growing degree days.

**Table S13. Total leaf numner from Iowa summer 2025 field.**

| Rep | GENOTYPE | P#1 | P#2 | P#3 | P#4 | P#5 | P#6 | P#7 | P#8 | P#9 | P#10 | P#11 | P#12 | P#13 | P#14 | P#15 | P#16 | P#17 | P#18 |
|---|---|---|---|---|---|---|---|---|---|---|---|---|---|---|---|---|---|---|---|
| **1** | 2X HO | 19 | 18 | 21 | 20 | | 20 | 19 | 19 | - | 20 | 20 | 19 | 19 | 20 | 19 | - | 20 | 20 |
| **2** | 2X HO | 20 | 19 | 20 | 19 | 19 | 20 | 21 | 21 | 20 | 19 | 20 | 20 | 21 | 21 | 21 | 21 | 20 | - |
| **3** | 2X HO | 20 | 20 | 20 | 20 | 20 | 20 | 20 | 20 | 20 | - | 20 | 20 | 20 | 20 | 21 | - | - | - |
| **4** | 2X HO | | 20 | 20 | 20 | 20 | 20 | 20 | 20 | 20 | 20 | 20 | - | - | 20 | 20 | | - | - |
| **5** | 2X HO | 24 | 24 | 21 | 24 | 20 | 20 | 20 | 20 | 20 | 21 | 19 | 20 | - | - | - | - | - | - |
| **1** | 3X HO | 18 | 19 | 19 | 19 | 20 | 20 | 20 | - | 19 | 20 | 19 | 19 | 18 | 20 | 20 | 20 | 20 | - |
| **2** | 3X HO | 20 | 20 | 19 | 20 | 20 | - | 21 | 20 | 20 | 20 | - | - | 20 | 20 | 19 | 18 | | 20 |
| **3** | 3X HO | 19 | 19 | 20 | 19 | 19 | 20 | 19 | 19 | 19 | 20 | 20 | 20 | 20 | 19 | - | - | - | - |
| **4** | 3X HO | | 20 | 21 | 19 | 20 | 19 | 18 | 19 | 19 | - | 19 | 20 | 18 | 19 | 21 | 19 | - | - |
| **5** | 3X HO | 20 | | 19 | 21 | 19 | 19 | 21 | 19 | 19 | 18 | 18 | 19 | 19 | 19 | 19 | 19 | - | 19 |
| **1** | Tzi8 | - | 24 | - | 22 | 22 | 23 | 21 | - | - | 22 | 23 | - | 24 | 24 | - | 23 | - | - |
| **2** | Tzi8 | - | 23 | - | - | - | 21 | - | - | 24 | - | - | - | - | 22 | - | - | - | - |
| **3** | Tzi8 | | 25 | 24 | 21 | - | 24 | - | - | 24 | 24 | - | - | 24 | 24 | 25 | 24 | - | - |
| **4** | Tzi8 | 22 | 22 | - | - | - | - | 22 | 22 | - | 23 | - | 21 | 24 | - | 21 | 24 | - | - |
| **5** | Tzi8 | - | - | - | - | 23 | 23 | - | 23 | 22 | 21 | 22 | 22 | 23 | 25 | - | - | - | - |

2X HO is the fixed double-edited *zmcct10, zmrap2.7* line.

3X HO is the fixed triple-edited *zmcct9, zmcct10, zmrap2.7* line.

**Table S14. Flowering traits from Iowa summer 2025 field.**

| Rep | GENOTYPE | Trait | P#1 | P#2 | P#3 | P#4 | P#5 | P#6 | P#7 | P#8 | P#9 | P#10 | P#11 | P#12 | P#13 | P#14 | P#15 | P#16 | P#17 | P#18 |
|---|---|---|---|---|---|---|---|---|---|---|---|---|---|---|---|---|---|---|---|---|
| 1 | 3X HO | DTA | 75 | 71 | 71 | - | 77 | 71 | 72 | 71 | 73 | 72 | 70 | 70 | 69 | 69 | 72 | 79 | 74 | |
| 2 | 3X HO | DTA | 72 | 72 | 70 | 72 | 74 | 75 | 74 | 70 | 77 | 72 | - | - | 78 | 73 | 75 | 81 | - | 73 |
| 3 | 3X HO | DTA | 80 | 78 | - | 78 | - | 78 | 72 | 77 | 81 | 77 | 81 | 75 | 77 | 79 | - | | | |
| 4 | 3X HO | DTA | - | 71 | 79 | 83 | 74 | 71 | 73 | 78 | - | - | 73 | 76 | 84 | 73 | - | 74 | | |
| 5 | 3X HO | DTA | 71 | - | - | 79 | 73 | 76 | 71 | 74 | 72 | 72 | - | 76 | 77 | 72 | 75 | 74 | - | 71 |
| 1 | 2X HO | DTA | 74 | - | 73 | - | - | 74 | - | - | - | 77 | 74 | 74 | - | - | - | - | 76 | 73 |
| 2 | 2X HO | DTA | 76 | 71 | 78 | 72 | 71 | 76 | 76 | 71 | 79 | 77 | 76 | 75 | 74 | 74 | 76 | 73 | 74 | |
| 3 | 2X HO | DTA | 74 | 72 | 78 | 73 | 74 | 73 | 77 | 73 | 76 | 77 | 74 | 74 | - | - | 77 | 83 | | |
| 4 | 2X HO | DTA | - | 82 | 74 | 79 | | 74 | - | 73 | 73 | 75 | 76 | - | 78 | 74 | 73 | | | |
| 5 | 2X HO | DTA | 72 | 70 | 73 | 71 | 71 | 71 | 71 | 74 | 71 | 71 | 71 | 71 | | | | | | |
| 1 | Tzi8 | DTA | - | - | - | 87 | 85 | | 81 | - | - | 74 | 93 | - | - | 83 | - | 80 | | |
| 2 | Tzi8 | DTA | - | 85 | - | - | - | 82 | - | - | 88 | - | - | - | - | 77 | - | | | |
| 3 | Tzi8 | DTA | - | - | 83 | - | - | 91 | - | - | 84 | 82 | - | - | 84 | - | 83 | - | | |
| 4 | Tzi8 | DTA | 79 | 84 | - | - | - | - | 78 | 85 | - | - | - | 77 | 84 | - | 74 | 87 | | |
| 5 | Tzi8 | DTA | - | - | - | - | 77 | 83 | - | - | 82 | 81 | 82 | - | 87 | - | | | | |
| 1 | 3X HO | DTS | 75 | 74 | 74 | 73 | 80 | 70 | 70 | - | 72 | 71 | 71 | 69 | 69 | 69 | 70 | 78 | 71 | |
| 2 | 3X HO | DTS | 75 | 76 | 71 | - | - | - | - | - | 79 | 76 | - | - | 79 | - | 75 | 79 | - | 74 |
| 3 | 3X HO | DTS | 74 | 78 | 73 | 77 | - | 76 | 73 | 73 | 77 | 75 | 82 | 73 | 76 | 72 | - | | | |
| 4 | 3X HO | DTS | - | 73 | 79 | 80 | 69 | 72 | - | 77 | 77 | - | 73 | 70 | 79 | 73 | 78 | 74 | | |
| 5 | 3X HO | DTS | 72 | - | 75 | 80 | 73 | 76 | 76 | 77 | 76 | 75 | 71 | 76 | 73 | 71 | 77 | 72 | - | 76 |
| 1 | 2X HO | DTS | - | 77 | 84 | 76 | | 73 | 77 | 72 | - | - | 71 | 71 | 73 | 72 | 76 | - | - | 73 |
| 2 | 2X HO | DTS | | 72 | 78 | 73 | 71 | 76 | - | 71 | 76 | 76 | 75 | 73 | 76 | 76 | 75 | 74 | 73 | |
| 3 | 2X HO | DTS | 75 | 72 | - | 79 | - | 73 | 75 | 73 | 75 | - | 76 | 74 | 79 | 73 | 70 | 82 | | |
| 4 | 2X HO | DTS | - | 82 | 74 | 78 | 78 | 72 | 81 | 75 | 77 | 75 | 75 | - | 76 | - | | | | |
| 5 | 2X HO | DTS | 75 | - | 76 | 72 | 72 | 70 | 73 | 75 | 73 | 72 | 73 | 72 | | | | | | |
| 1 | Tzi8 | DTS | 83 | - | 82 | 86 | 82 | 82 | 80 | - | 86 | 76 | 79 | - | 82 | 82 | 91 | 80 | | |
| 2 | Tzi8 | DTS | 82 | 85 | 79 | 80 | 82 | 79 | 81 | 82 | - | - | 83 | - | 81 | 76 | 85 | | | |
| 3 | Tzi8 | DTS | - | 82 | 82 | 80 | 91 | 85 | 82 | 85 | 78 | - | 88 | 81 | 80 | 82 | 82 | 82 | | |
| 4 | Tzi8 | DTS | 78 | 82 | 87 | - | 81 | - | 79 | 87 | 83 | 85 | 82 | 77 | 79 | 88 | 75 | 80 | | |
| 5 | Tzi8 | DTS | 86 | 79 | 91 | 81 | 77 | 82 | - | 81 | 77 | 80 | - | 78 | 82 | 81 | | | | |
| 1 | 3X HO | GDD_DTA | 1806.5 | 1712.5 | 1712.5 | - | 1861.5 | 1712.5 | 1735 | 1712.5 | 1760.5 | 1735 | 1688 | 1688 | 1660 | 1660 | 1735 | 1918 | 1782.5 | |
| 2 | 3X HO | GDD_DTA | 1735 | 1735 | 1688 | 1735 | 1782.5 | 1806.5 | 1782.5 | 1688 | 1861.5 | 1735 | - | - | 1890.5 | 1760.5 | 1806.5 | 1973 | - | 1760.5 |
| 3 | 3X HO | GDD_DTA | 1945.5 | 1890.5 | - | 1890.5 | - | 1890.5 | 1735 | 1861.5 | 1973 | 1861.5 | 1973 | 1806.5 | 1861.5 | 1918 | - | | | |
| 4 | 3X HO | GDD_DTA | - | 1712.5 | 1918 | 2021 | 1782.5 | 1712.5 | 1760.5 | 1890.5 | - | - | 1760.5 | 1833 | 2042 | 1760.5 | - | 1782.5 | | |
| 5 | 3X HO | GDD_DTA | 1712.5 | - | - | 1918 | 1760.5 | 1833 | 1712.5 | 1782.5 | 1735 | 1735 | - | 1833 | 1861.5 | 1735 | 1806.5 | 1782.5 | - | 1712.5 |
| 1 | 2X HO | GDD_DTA | 1782.5 | - | 1760.5 | - | - | 1782.5 | - | - | - | 1861.5 | 1782.5 | 1782.5 | - | - | - | - | 1833 | 1760.5 |
| 2 | 2X HO | GDD_DTA | 1833 | 1712.5 | 1890.5 | 1735 | 1712.5 | 1833 | 1833 | 1712.5 | 1918 | 1861.5 | 1833 | 1806.5 | 1782.5 | 1782.5 | 1833 | 1760.5 | 1782.5 | |
| 3 | 2X HO | GDD_DTA | 1782.5 | 1735 | 1890.5 | 1760.5 | 1782.5 | 1760.5 | 1861.5 | 1760.5 | 1833 | 1861.5 | 1782.5 | 1782.5 | - | - | 1861.5 | 2021 | | |
| 4 | 2X HO | GDD_DTA | - | 1998 | 1782.5 | 1918 | | 1782.5 | - | 1760.5 | 1760.5 | 1806.5 | 1833 | - | 1890.5 | 1782.5 | 1760.5 | | | |
| 5 | 2X HO | GDD_DTA | 1735 | 1688 | 1760.5 | 1712.5 | 1712.5 | 1712.5 | 1712.5 | 1782.5 | 1712.5 | 1712.5 | 1712.5 | 1712.5 | | | | | | |
| 1 | Tzi8 | GDD_DTA | - | - | - | 2086.5 | 2061 | | 1973 | - | - | 1782.5 | 2205.5 | - | - | 2021 | - | 1945.5 | | |
| 2 | Tzi8 | GDD_DTA | - | 2061 | - | - | - | 1998 | - | - | 2104 | - | - | - | - | 1861.5 | - | | | |
| 3 | Tzi8 | GDD_DTA | - | - | 2021 | - | - | 2164 | - | - | 2042 | 1998 | - | - | 2042 | - | 2021 | - | | |
| 4 | Tzi8 | GDD_DTA | 1918 | 2042 | - | - | - | - | 1890.5 | 2061 | - | - | - | 1861.5 | 2042 | - | 1782.5 | 2086.5 | | |
| 5 | Tzi8 | GDD_DTA | - | - | - | - | 1861.5 | 2021 | - | - | 1998 | 1973 | 1998 | - | 2086.5 | - | | | | |
| 1 | 3X HO | GDD_DTS | 1806.5 | 1782.5 | 1782.5 | 1760.5 | 1945.5 | 1688 | 1688 | - | 1735 | 1712.5 | 1712.5 | 1660 | 1660 | 1660 | 1688 | 1890.5 | 1712.5 | |
| 2 | 3X HO | GDD_DTS | 1806.5 | 1833 | 1712.5 | - | - | - | - | - | 1918 | 1833 | - | - | 1918 | - | 1806.5 | 1918 | - | 1782.5 |
| 3 | 3X HO | GDD_DTS | 1782.5 | 1890.5 | 1760.5 | 1861.5 | - | 1833 | 1760.5 | 1760.5 | 1861.5 | 1806.5 | 1998 | 1760.5 | 1833 | 1735 | - | | | |

| | | | | | | | | | | | | | | | | | | | | |
|---|---|---|---|---|---|---|---|---|---|---|---|---|---|---|---|---|---|---|---|---|
| **4** | 3X HO | GDD_DTS | - | 1760.5 | 1918 | 1945.5 | 1660 | 1735 | - | 1861.5 | 1861.5 | - | 1760.5 | 1688 | 1918 | 1760.5 | 1890.5 | 1782.5 | | |
| **5** | 3X HO | GDD_DTS | 1735 | - | 1806.5 | 1945.5 | 1760.5 | 1833 | 1833 | 1861.5 | 1833 | 1806.5 | 1712.5 | 1833 | 1760.5 | 1712.5 | 1861.5 | 1735 | - | 1833 |
| **1** | 2X HO | GDD_DTS | - | 1861.5 | 2042 | 1833 | | 1760.5 | 1861.5 | 1735 | - | - | 1712.5 | 1712.5 | 1760.5 | 1735 | 1833 | - | - | 1760.5 |
| **2** | 2X HO | GDD_DTS | | 1735 | 1890.5 | 1760.5 | 1712.5 | 1833 | - | 1712.5 | 1833 | 1833 | 1806.5 | 1760.5 | 1833 | 1833 | 1806.5 | 1782.5 | 1760.5 | |
| **3** | 2X HO | GDD_DTS | 1806.5 | 1735 | - | 1918 | - | 1760.5 | 1806.5 | 1760.5 | 1806.5 | - | 1833 | 1782.5 | 1918 | 1760.5 | 1688 | 1998 | | |
| **4** | 2X HO | GDD_DTS | - | 1998 | 1782.5 | 1890.5 | 1890.5 | 1735 | 1973 | 1806.5 | 1861.5 | 1806.5 | 1806.5 | - | 1833 | - | | | | |
| **5** | 2X HO | GDD_DTS | 1806.5 | - | 1833 | 1735 | 1735 | 1688 | 1760.5 | 1806.5 | 1760.5 | 1735 | 1760.5 | 1735 | | | | | | |
| **1** | Tzi8 | GDD_DTS | 2021 | - | 1998 | 2073 | 1998 | 1998 | 1945.5 | - | 2073 | 1833 | 1918 | - | 1998 | 1998 | 2164 | 1945.5 | | |
| **2** | Tzi8 | GDD_DTS | 1998 | 2061 | 1918 | 1945.5 | 1998 | 1918 | 1973 | 1998 | - | - | 2021 | - | 1973 | 1833 | 2061 | | | |
| **3** | Tzi8 | GDD_DTS | - | 1998 | 1998 | 1945.5 | 2164 | 2061 | 1998 | 2061 | 1890.5 | - | 2104 | 1973 | 1945.5 | 1998 | 1998 | 1998 | | |
| **4** | Tzi8 | GDD_DTS | 1890.5 | 1998 | 2086.5 | - | 1973 | - | 1918 | 2086.5 | 2021 | 2061 | 1998 | 1861.5 | 1918 | 2104 | 1806.5 | 1945.5 | | |
| **5** | Tzi8 | GDD_DTS | 2073 | 1918 | 2164 | 1973 | 1861.5 | 1998 | - | 1973 | 1861.5 | 1945.5 | - | 1890.5 | 1998 | 1973 | | | | |

2X HO is the fixed double-edited *zmcct10, zmrap2.7* line.

3X HO is the fixed triple-edited *zmcct9, zmcct10, zmrap2.7* line.

DTA, days to anthesis; DTS, days to silking, GDD, growing degree days.
